## Supplementary Figures for "Integrating Cell Morphology with Gene Expression and Chemical Structure to Aid Mitochondrial Toxicity Detection"

Mitochondrial Toxicity; Cell Painting; Toxicity Prediction; Cell Morphology

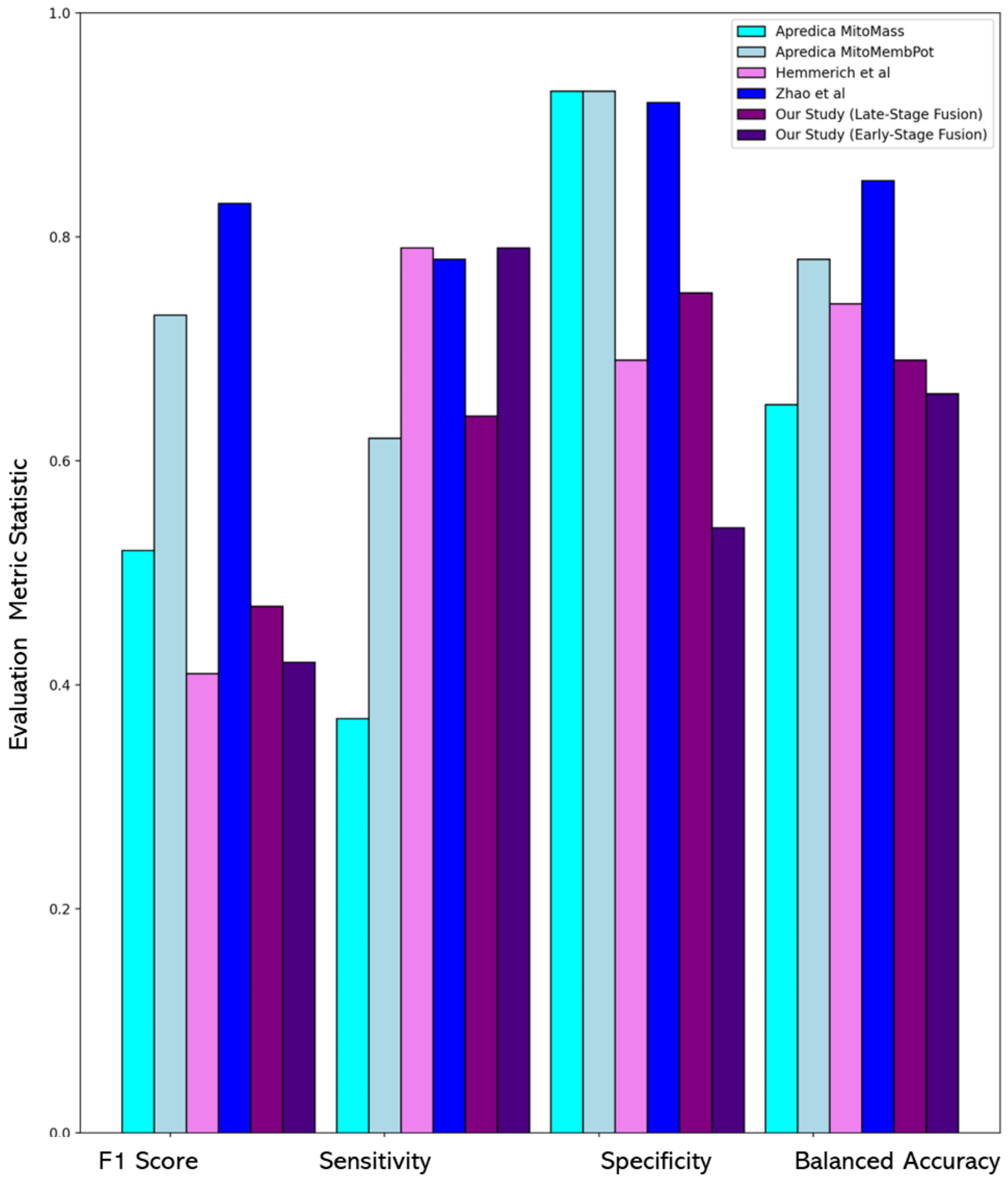


**Figure S1.** Comparison of our models with models previously published and dedicated high content imaging assays for mitochondrial toxicity. Although the test set compounds differ and results are not directly comparable. The early-stage fusion models improved F1 Scores compared to Hemmerich et al.[1] (0.47 vs 0.41) Our late-stage fusion model achieve comparable balanced accuracy to previous computational models by Hemmerich et al.[1] (0.69 vs 0.74) and better than two of the high content imaging assays (0.79 vs 0.62 in Apredica MitoMembPot and 0.37 in Apredica MitoMass).

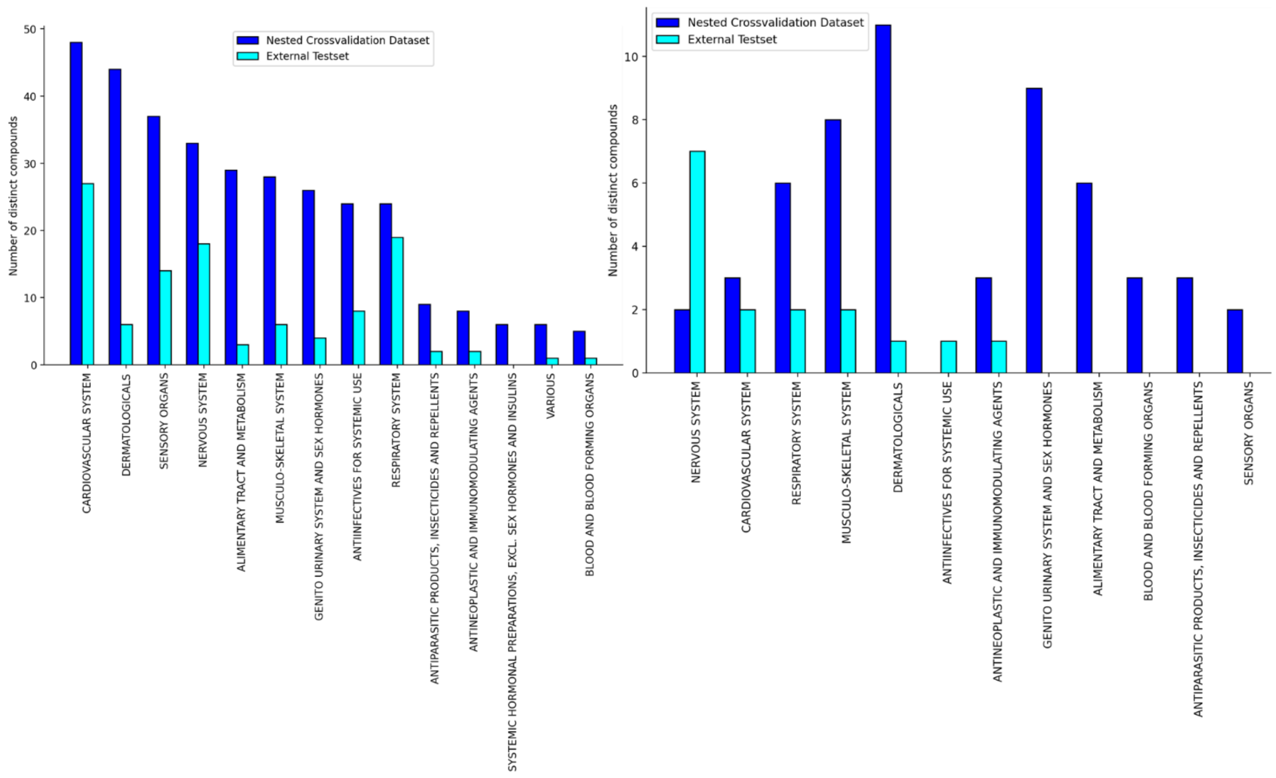


**Figure S2.** (a) Empirical ATC distribution at the top level for 327 drugs out of 382 compounds in the training data and 111 drugs out of 244 compounds in the external test set. (b) Empirical ATC distribution at the top level for 56 mitochondrial toxicants out of 62 toxicants in the training data and 16 mitochondrial toxicants out of 47 toxicants in the external test set.

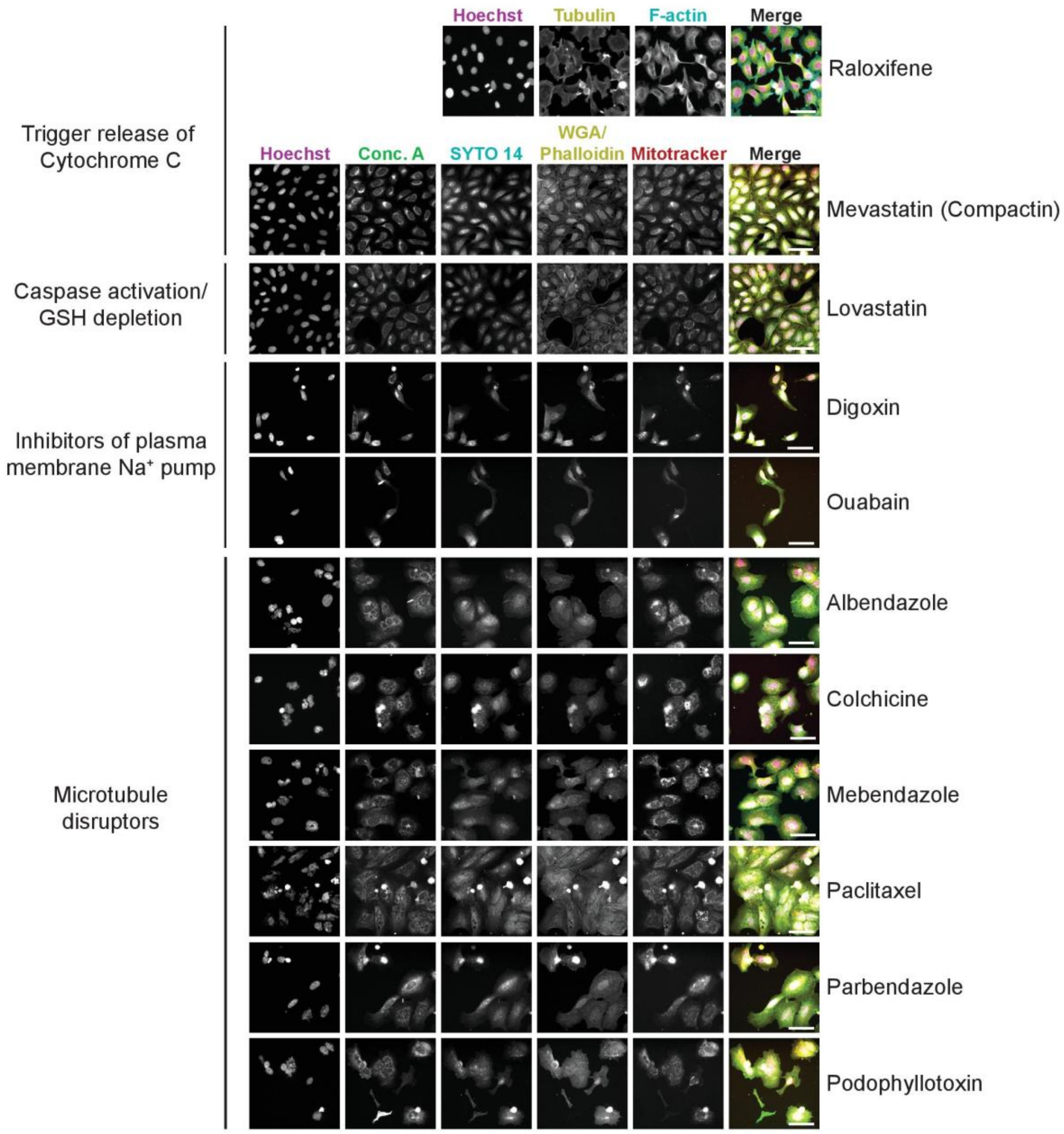


**Figure S3.** Representative images of cells stained using the Cell Painting assay upon exposed to drugs with different mode of action.

**Raloxifene** is an estrogen modulator and **Mevastatin** is cholesterol-lowering drug which inhibits the HMG-Coenzyme A reductase. Both drugs are found to induce the release of cytochrome C in the mitochondria. Similarly to Mevastatin, **Lovastatin** is cholesterol-lowering drug which also inhibits the HMG-Coenzyme A reductase, however it has been found to induce apoptosis, G1 phase cell cycle arrest and GSH depletion. **Digoxin** is a cardiac glycoside that inhibits sodium potassium ATPase pumps, and it has found to induce apoptosis; **Ouabain** is a steroid hormone that also inhibits sodium potassium ATPase pumps. The cell painting phenotype reveals a clear toxic effect on U2OS cells, depicted by the low number of remaining cells after exposure. **Albendazole**, **Colchicine**, **Mebendazole**, **Paclitaxel**, **Parbendazole** and **Podophyllotoxin** are six microtubule disruptor drugs that have been found to induce cytotoxicity. The cell painting phenotype reveals alterations at the nuclear level, depicted by nuclear fragmentation as well as multinucleated cells, vacuolation of the endoplasmic reticulum, redistribution of the mitochondria and cytoskeleton destabilisation.

These images are publicly available through the Broad Bioimage Benchmark Collection (https://bbbc.broadinstitute.org/image_sets), particularly the image set BBBC021 (Human MCF7 cells – compound-profiling experiment) from which images of MCF7 cells exposed to **Raloxifene** were extracted; and BBBC022 (Human U2OS cells – compound-profiling Cell Painting experiment) from which images of U2OS cells exposed to **Mevastatin**, **Lovastatin**, **Digoxin**, **Ouabain**, **Albendazole**, **Colchicine**, **Mebendazole**, **Paclitaxel**, **Parbendazole** and **Podophyllotoxin** were extracted. The scale bar, 50 µm (indicated in the merged images).

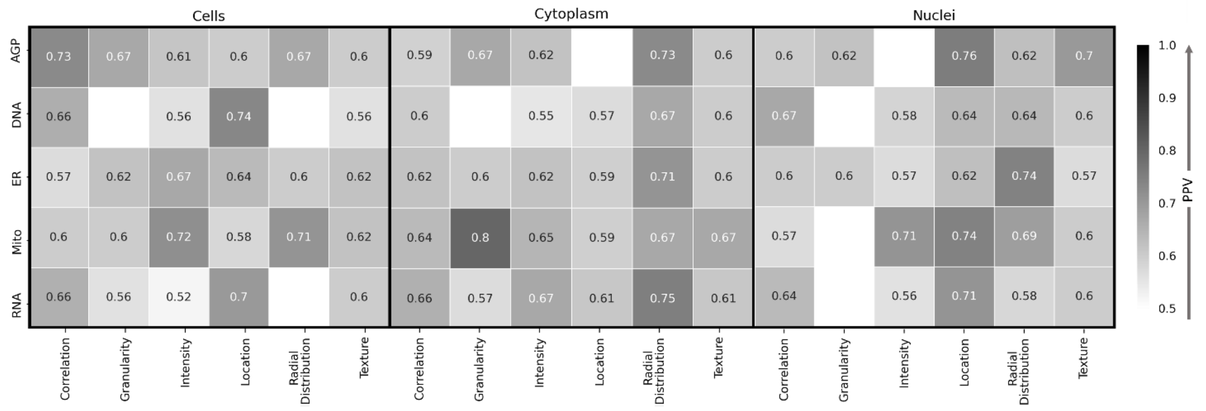


**Figure S4.** The median Positive Predictive Value (PPV) of Cell Painting features having PPV>0 in predicting mitochondrial toxicity from 486 compounds. The features were grouped by compartment (Cells, Cytoplasm, and Nuclei), channel (AGP, Nucleus, ER, Mito, Nucleolus/Cyto RNA), and feature group (Correlation, Granularity, Intensity, Radial Distribution, Texture). White squares indicate that no features with given combination had a PPV>0. Granularity, intensity, location, and radial distribution of mitochondrial objects over the three compartments had high PPV (>=0.70).

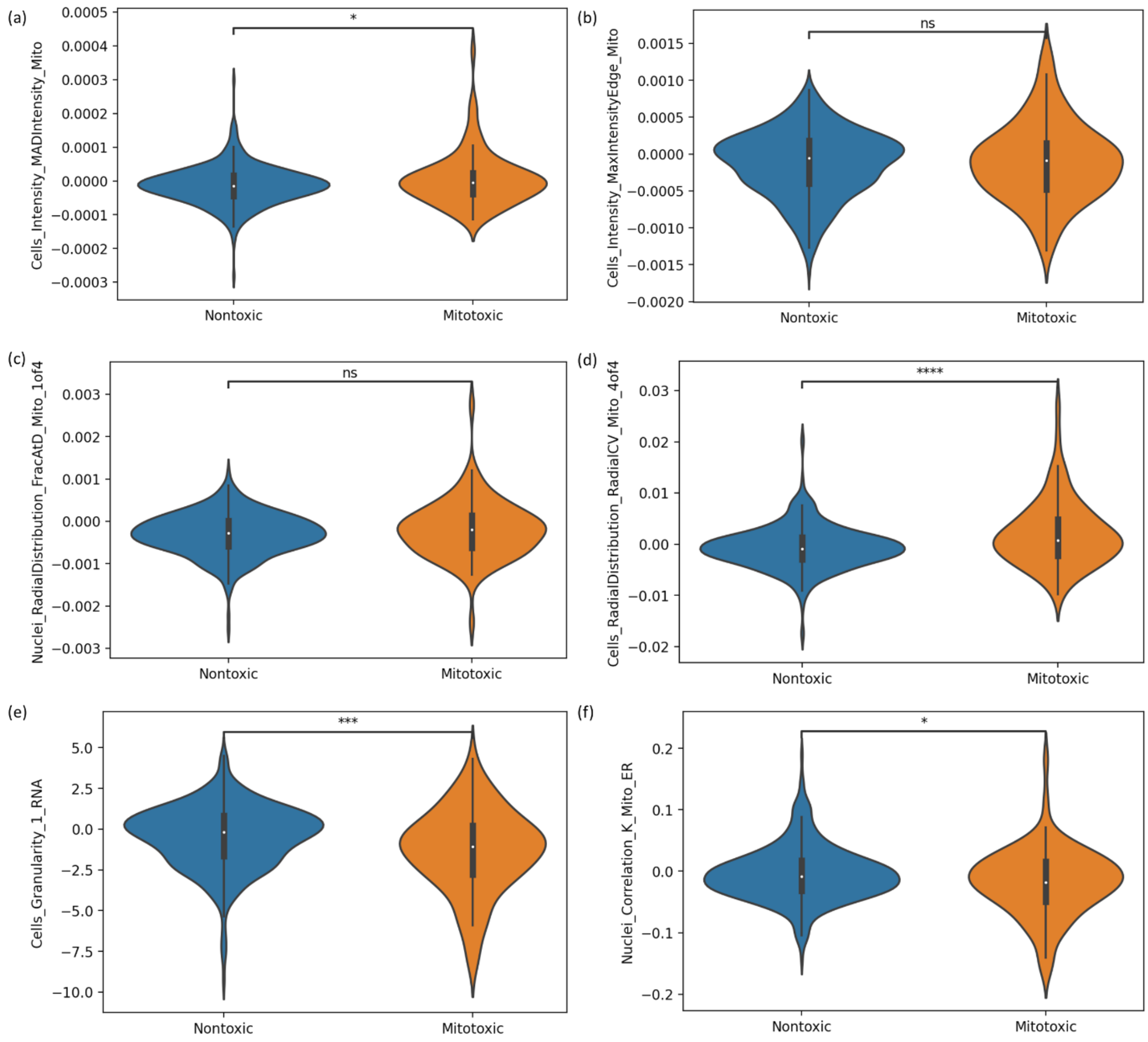


**Figure S5.** T-test independent samples with Bonferroni correction on distribution of selected individual Cell Painting feature for mitochondrial toxicants and non-toxic compounds.

(a) "Cells Intensity MADIntensity Mito" (PPV=0.8) is a measurement of statistical dispersion which measures the standard deviation and median absolute deviation (MAD) of pixel intensity values while being robust to outliers. For MitoTracker Deep Red used in Cell Painting assay, cells with reduced mitochondrial membrane potential will have less fluoresce and hence less intensity. For cell images of mitotoxic compounds, MADIntensity will have a higher positive value compared to non-toxic compounds which behave similarly to neutral controls, (this their difference is zero).

(b) “Cells Intensity MaxIntensityEdge Mito” (PPV=0.83) measures the maximal edge pixel intensity of an object, in our case the mitochondria which mean mitochondrial toxicants affect edges of mitochondria.

(c) “Nuclei RadialDistribution FracAtD Mito 1of4” (PPV=0.80) , the feature related to the innermost ring intensity of mitochondria. If the fraction of total stain mitochondria is lower in mitotoxic compounds, this indicates a loss of mitochondrial intensity, also in the proximity with the nucleus (since this is measured on the nuclei segmentation)

(d) “Cells RadialDistribution RadialCV Mito 4of4” (PPV=0.75) describes the coefficient of variation of intensity within the outer ring, calculated across 8 slices on the object (mitochondria) and was found to be higher in value for mitotoxic compounds compared to non-toxic compounds.

(e) “Cells Granularity 1 RNA” (PPV=0.56) reveals information present of pixel 1 in the RNA channel where certain mitotoxic compounds also have lower feature values. As this information is lower than the control after cell death, the feature value is more negative for compounds showing cytotoxicity. Most mitotoxic compounds have a value close to zero, indicating mitochondrial toxicity did not cause excessive cell death at the corresponding dose and time point in the Cell Painting assay.

(f) “Nuclei Correlation K Mito ER” (PPV=0.71), which calculates the correlation between intensities in different images on a pixel-by-pixel basis within mitochondria and ER. The contact between mitochondria and the ER can regulate biological processes such as ATP generation and mitochondrial division.

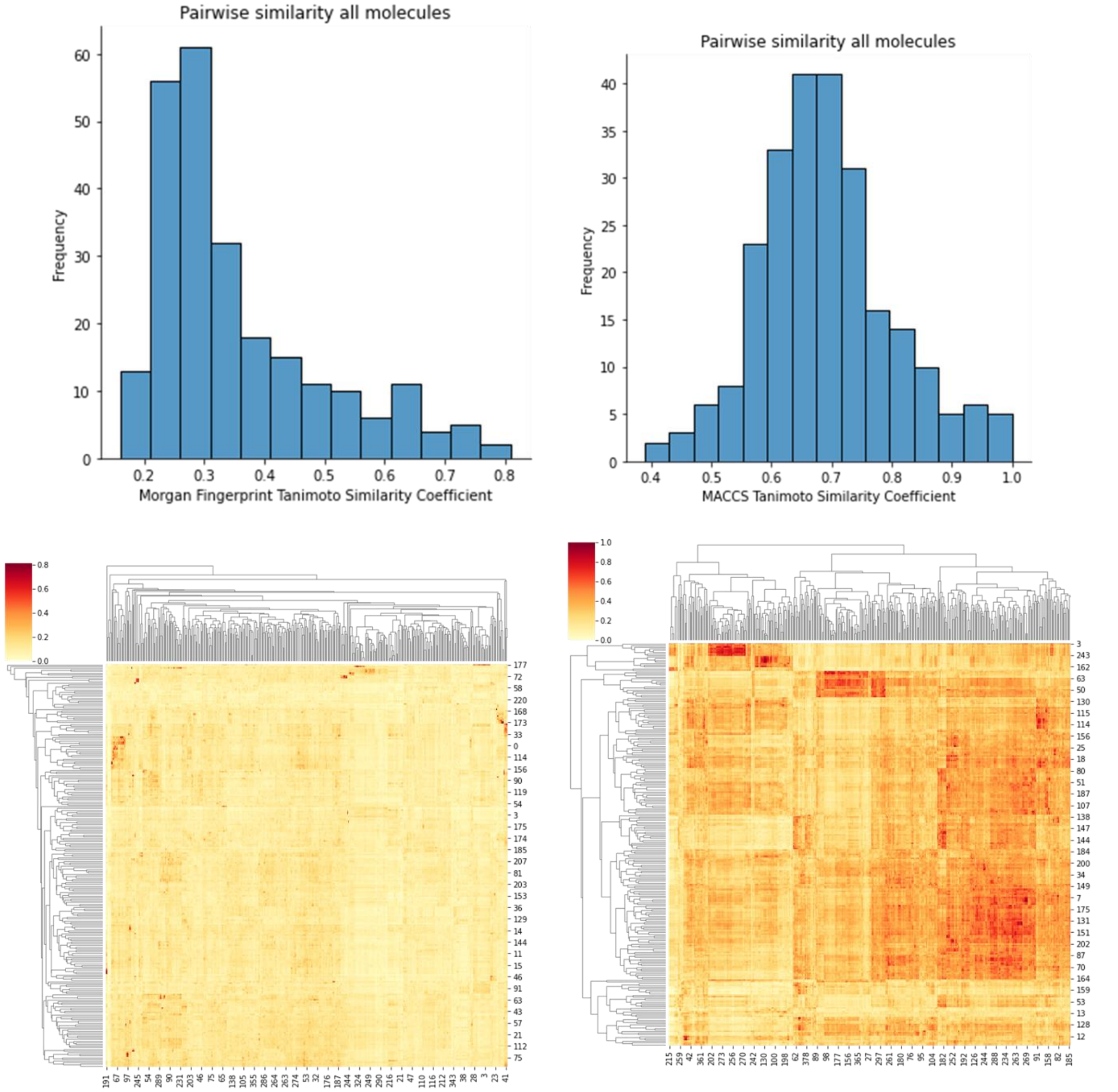


**Figure S6.** Pairwise combinations of training (382 compounds) vs test set (244 compounds) having highest Tanimoto similarity (Tc) using (a) Morgan Fingerprints of radius 2 and 2048 bits and (b) 166 public MACCS keys. (c) and (d) show clustered heatmaps of all pairwise Tanimoto similarity coefficients using Morgan Fingerprints of radius 2 and 2048 bits and 166 public MACCS keys respectively. This shows most test compounds are structurally diverse to the training set (with Tc<0.40 for Morgan Fingerprints and Tc<0.85 for MACCS keys).

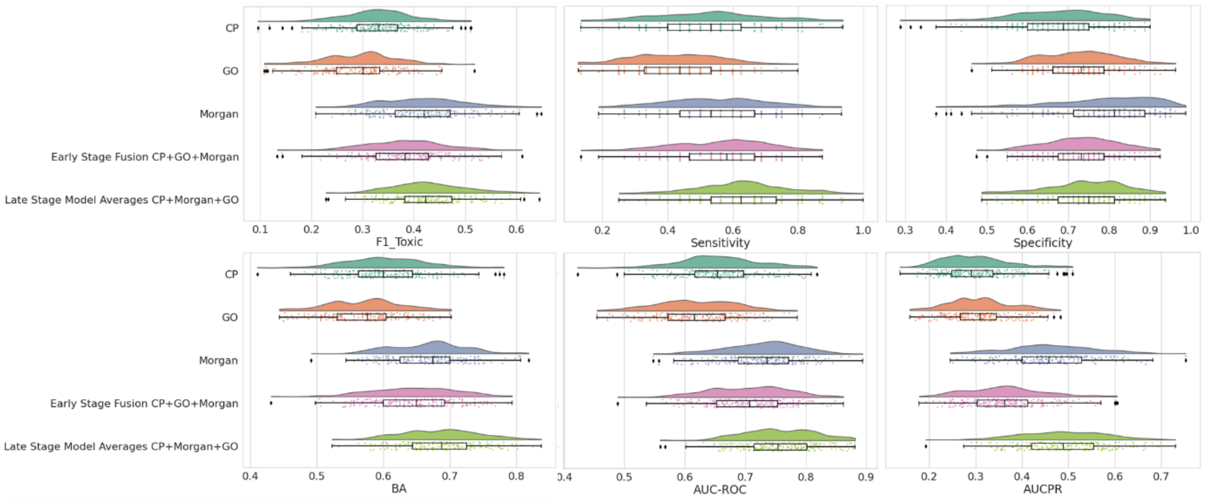


**Figure S7.** Evaluation metrics (F1 Score, Sensitivity, Specificity, Balanced Accuracy, AUC-ROC and AUCPR) for the five models for internal test sets from repeated nested cross validations. Late-stage models (median Sensitivity= 0.63, median AUC-ROC= 0.75, median AUC-PR= 0.49) recorded a relative improvement in sensitivity (by 17.2%), AUR-ROC (by 2.5%) and AUC-PR (by 6.7%) compared to Morgan fingerprints (median Sensitivity= 0.53, median AUC-ROC= 0.74, median AUC-PR= 0.46). CP: Cell Painting, GE: Gene Expression, BA: Balanced Accuracy

When considering results from 200 test sets from the nested cross-validation, Morgan fingerprints (median F1 score of 0.42 and median precision of 0.35) perform similar to late-stage fusion combining Cell Painting, Gene Expression and Morgan Fingerprints (median F1 score of 0.42 and median precision of 0.32). However, compared to sensitivity of 0.53 when using Morgan fingerprints, the late-stage fusion combining Cell Painting, Gene Expression and Morgan Fingerprints recorded a median sensitivity of 0.63 (relatively increased by 17.2%, T-test p-value=1.5e-11). The corresponding median specificity decreased from 0.81 when Morgan fingerprints to 0.75 when using late-stage fusion combining Cell Painting, Gene Expression and Morgan Fingerprints (relatively decreased by 7.7%, T-test p-value=2.2e-05). When considering AUR-ROC, late-stage fusion combining Cell Painting, Gene Expression and Morgan Fingerprints (median AUC-ROC= 0.75) was significantly improved by 2.5% (-test p-value=1.8e-09) compared to Morgan fingerprints (median AUC-ROC= 0.74). When considering mitotoxic classes only, AUC-PR for late-stage fusion combining Cell Painting, Gene Expression and Morgan Fingerprints (median AUC-PR =

0.49) was significantly improved by 6.7% (T-test p-value=6.8e-06) compared to Morgan fingerprints (median AUC-PR = 0.36).

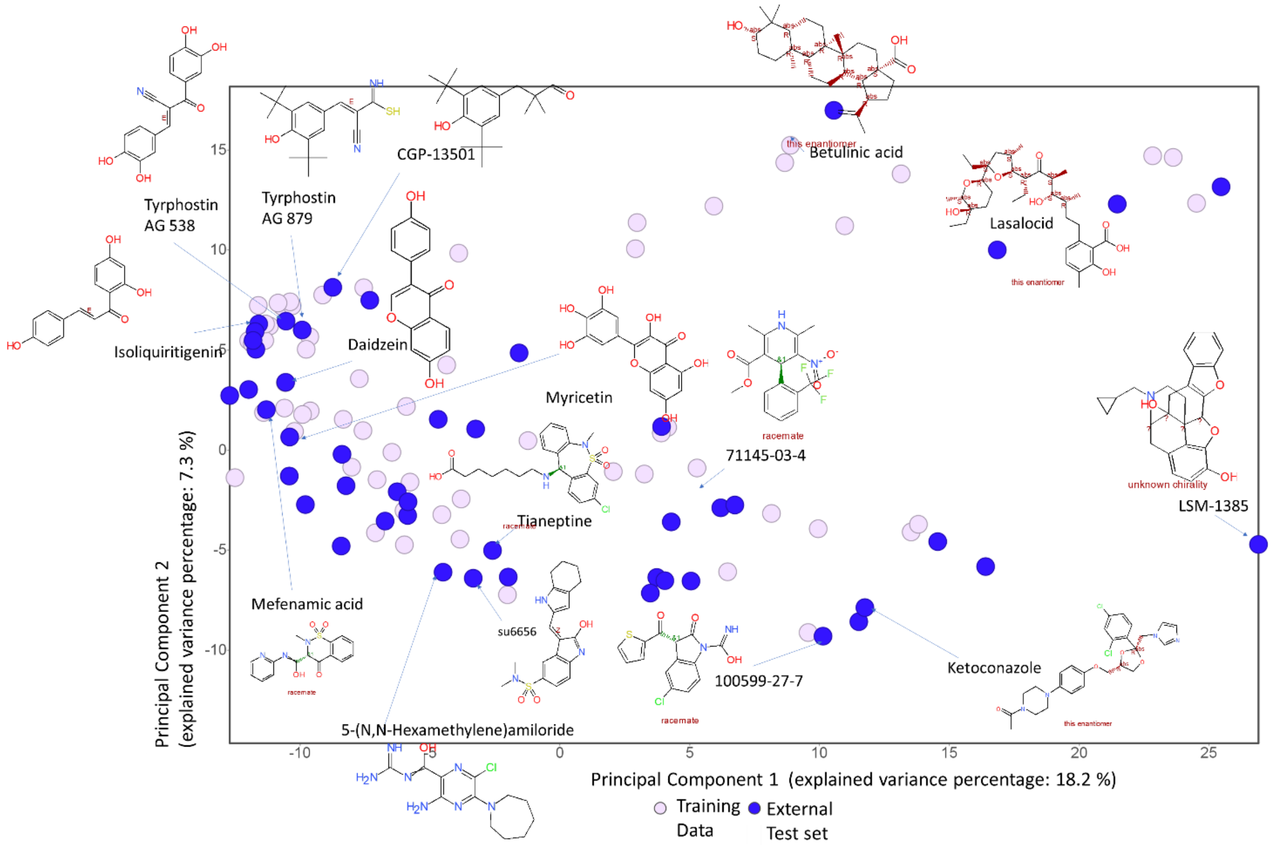


**Figure S8.** Mitochondrial toxicants in testing set compared to the training set in structural space defined by principal component analysis of FragFP fingerprints (DataWarrior).

Among mitochondrial toxicants in testing set, myricetin, mefenamic acid, betulinic acid and ketoconazole have been shown to inhibit the oxidative phosphorylation causing oxidative stress, [2,3,4] isoliquiritigenin and daidzein induces a ROS-mediated mitochondrial dysfunction pathway, [5] 5-(N,N-Hexamethylene)amiloride induces ROS generation[6], tyrphostin AG 879 induces oxidative stress[7], fluoxetine inhibits oxygen consumption and lowers mitochondrial ATP[8], su6656 activates AMP-activated protein kinase[9] and lasalocid increases Ca2+ influx[10], all causing the outer mitochondrial membrane to depolarize.

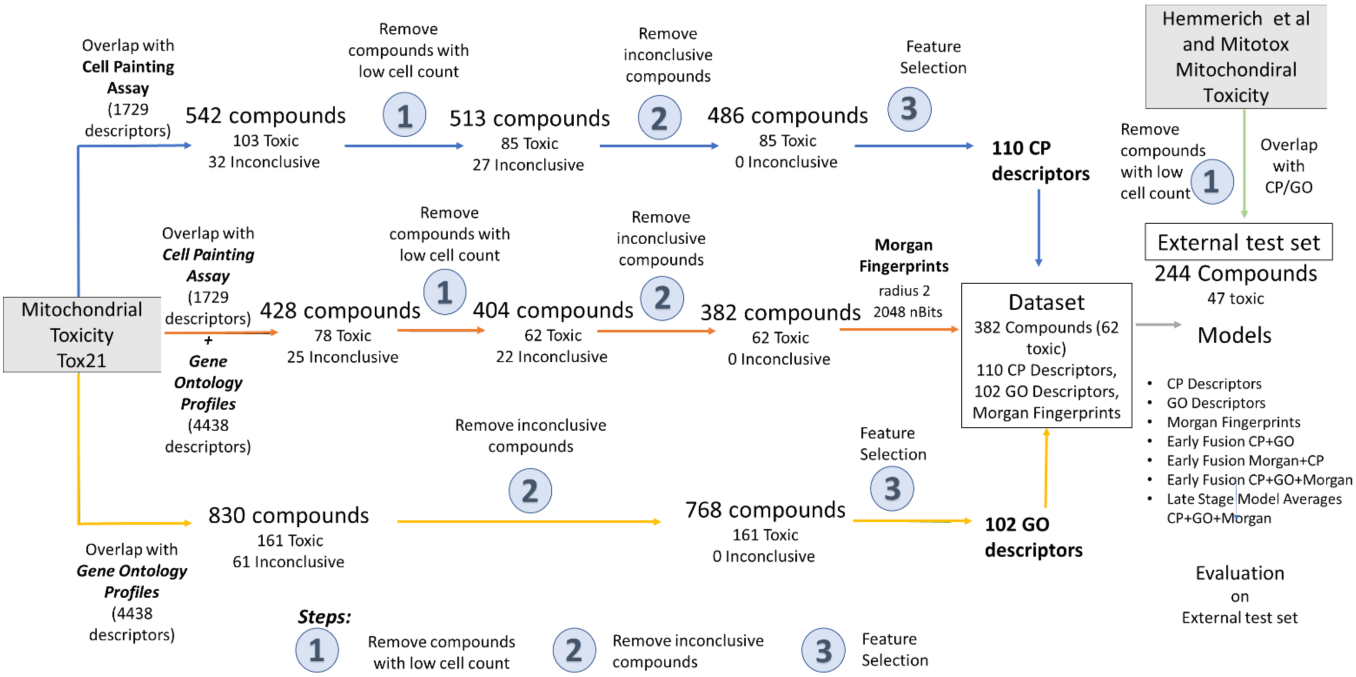


(a)

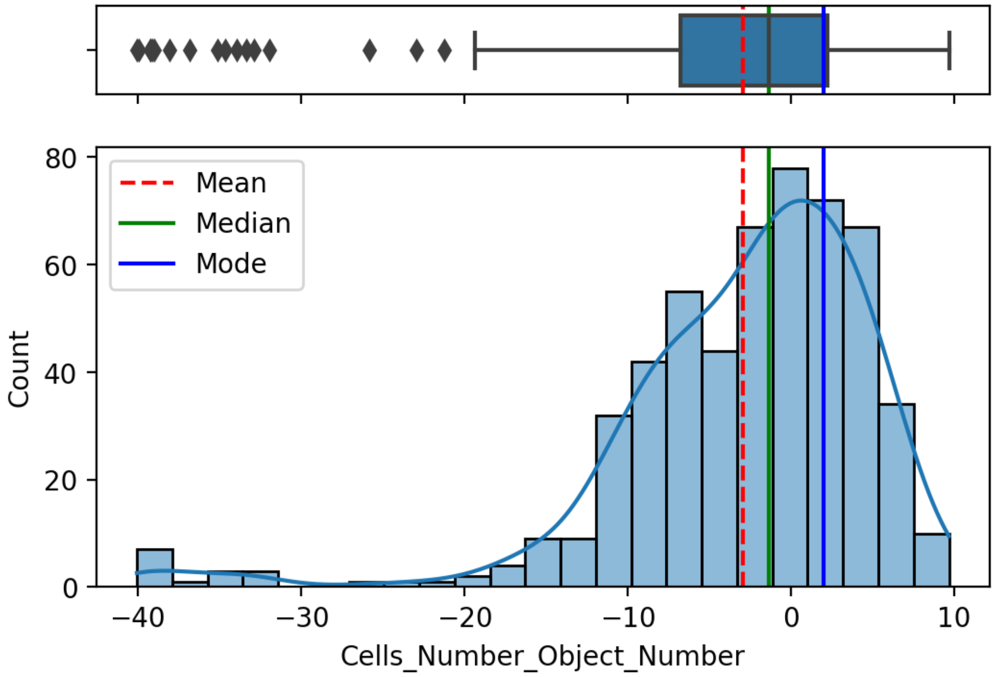


(b)

**Figure S9.** (a) Data collation workflow. Overlaps of mitotoxicity data were calculated with Cell Painting and Gene Expression data. When using Cell Painting features, we avoided compounds having low cell count (step 1, further in (b)) After removing the inconclusive compounds (step 2), the feature selection (step 3) was performed on Cell Painting features and Gene Expression annotations for the remaining compounds. The overlap of external test set was merged with available annotations of Cell Painting features and Gene Expression features were calculated in a similar manner, compounds with low cell count removed and no compounds from this was used in feature selection or training our models. (b) The distribution of Cell Count among the 542 compounds for which Cell Painting and Tox21 Mitochondrial toxicity annotations were available. We removed compounds which drastically reduced the cell count compared to the neutral control of DMSO (by a threshold of 1.5 times standard deviation below the mean of "Cells Number Object Number" which is below -15.09).

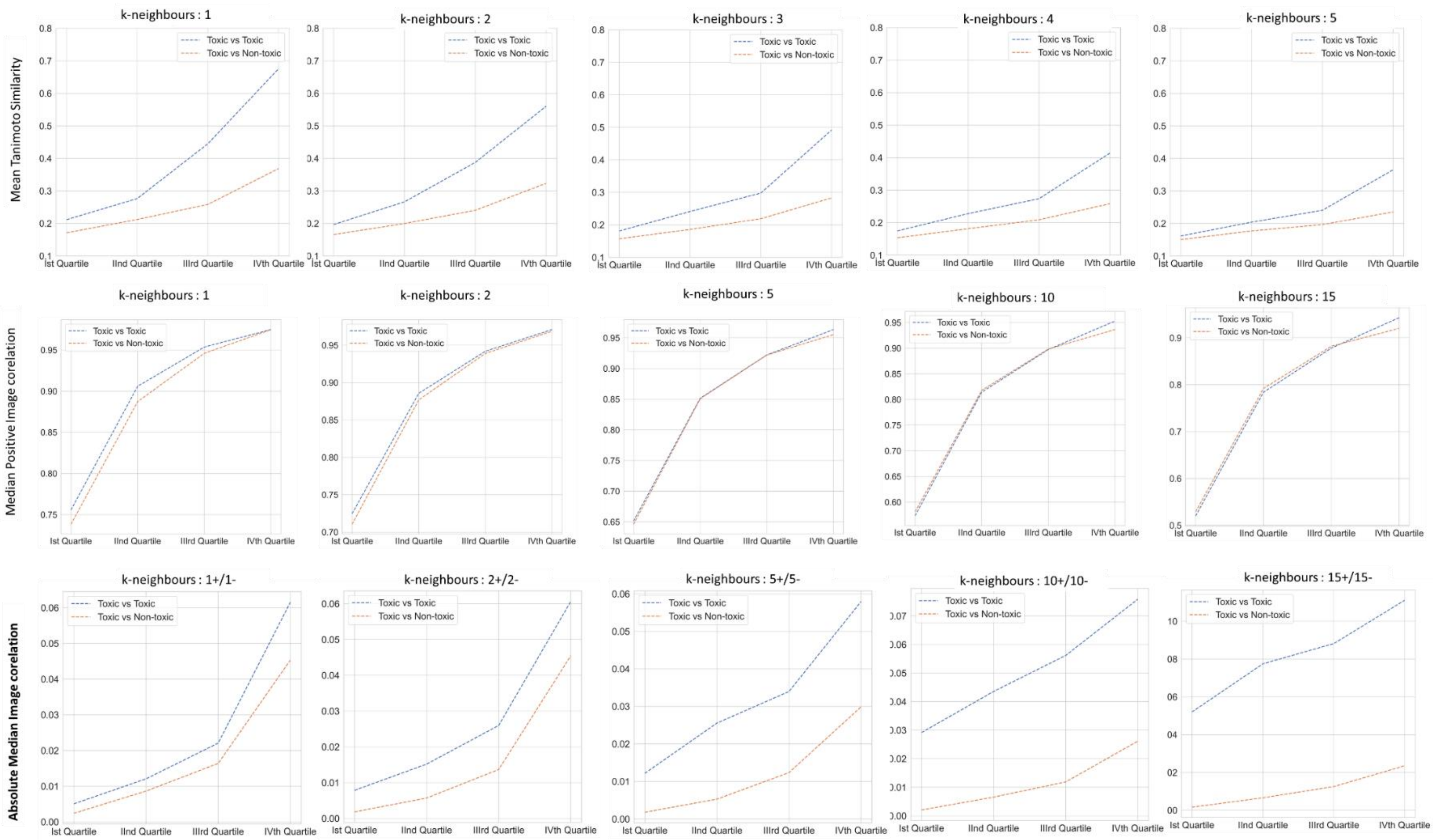


**Figure S10.** Mean Tanimoto Similarity, Median Positive Image Correlation and Absolute Median Image Correlation for k nearest neighbours in four quartiles of the distribution for Intra- and inter-class pairwise distributions for 486 compounds (85 mitotoxic). Better separation is obtained in using 5 neighbours for structural distance. For morphological distance, both positive and negative correlations for 15 neighbours give best separation of intra and inter class; hence this method was used to define the structural and morphological distance between training and test dataset as well.

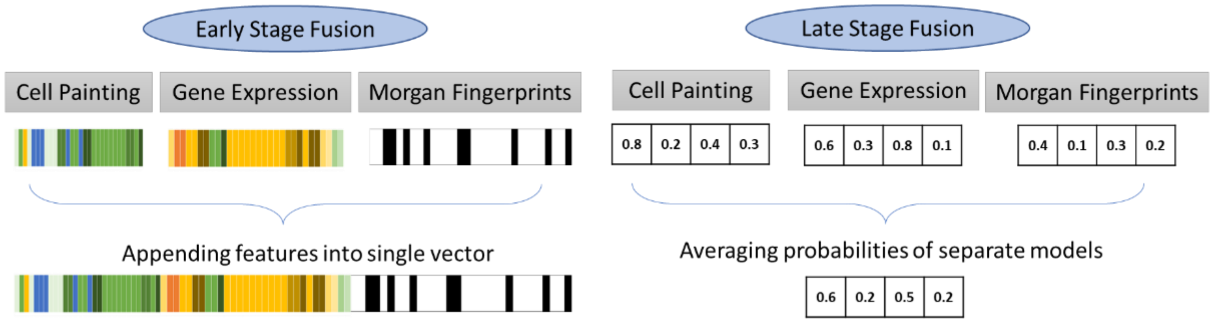


**Figure S11.** Fusion models were trained either by appending features together (early-stage) or averaging random forest prediction probabilities for the individual models (late-stage)

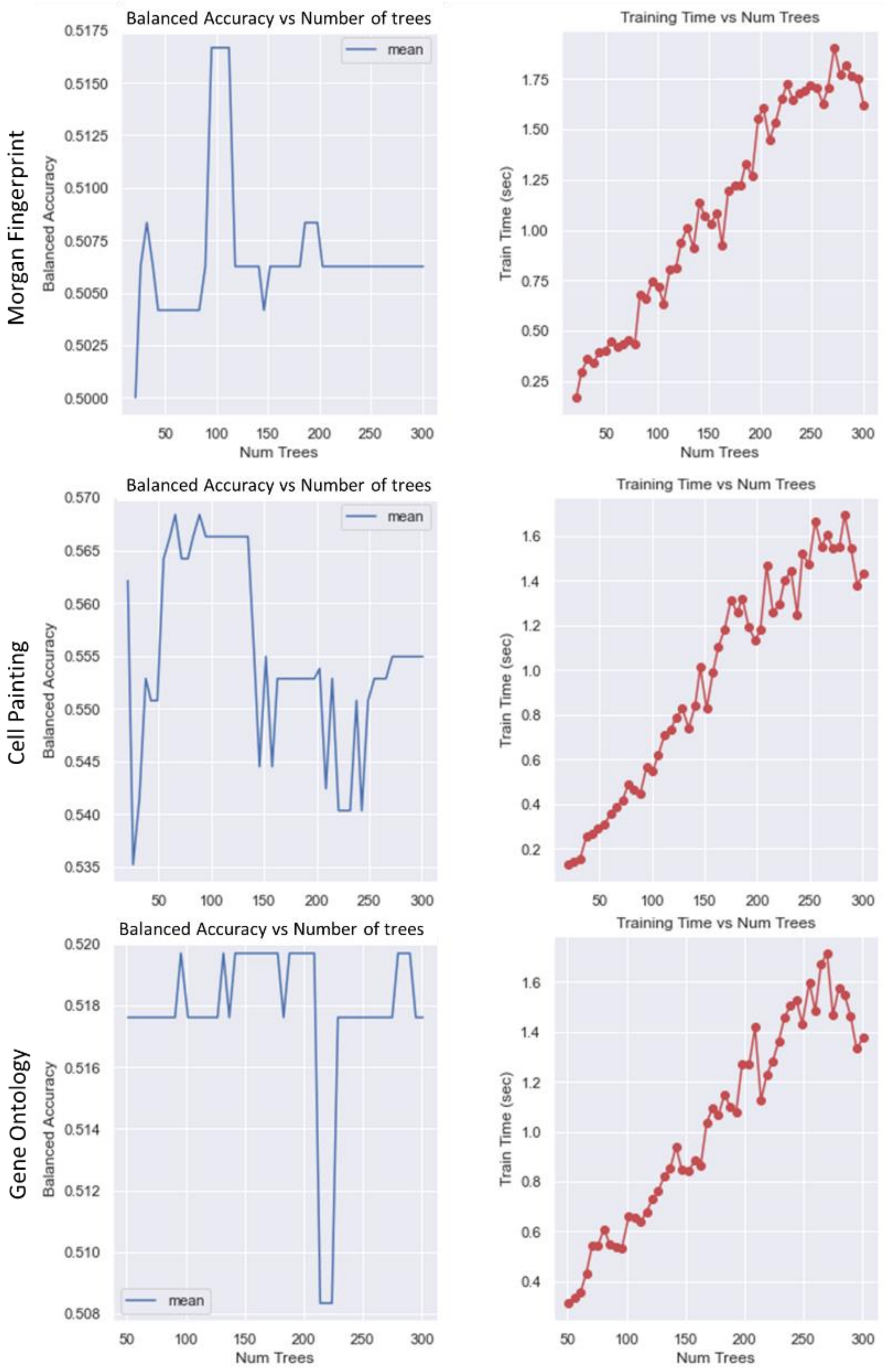


**Figure S12.** Hyperparameter tuning using grid search and a 4-fold stratified cross validation to determine optimal number of trees in random forest; our analysis revealed that 100 trees were capable in modelling using all three feature sets: Morgan fingerprints, Cell Painting and Gene Expression with reasonable training times. The consistent performance is most likely as random forests are usually robust against overfitting.

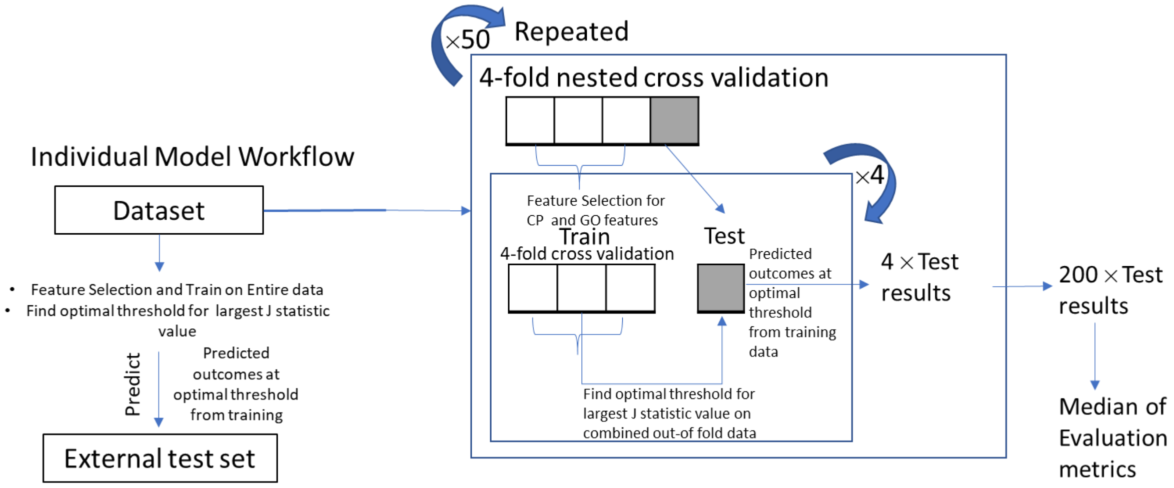


**Figure S13.** Modelling workflow used a 4-foldcross validation which allowed to vary the fourth fold of the data as internal test set such all the training data was included in the internal test set once. Feature selection for nested cross validation was performed only on the respective training data. This four-fold nested cross validation was repeated 50 times. The entire Model workflow was repeated for each model to analyse which feature/combinations are predictive of Mitochondrial toxicity. The external test predictions were made from models trained on the entire training set.
